# First biodegradable bioactive glass-based humidity sensor

**DOI:** 10.1101/2022.12.06.519262

**Authors:** Amina Gharbi, Ahmed Yahia Kallel, Olfa Kanoun, Wissem Cheikhrouhou-Koubaa, Christopher H. Contag, Iulian Antoniac, Nabil Derbel, Nureddin Ashammakhi

## Abstract

Monitoring changes in edema-associated intracranial pressure that complicates trauma or surgery would lead to improved outcomes. Implantable pressure sensors have been explored, but, these sensors require post-surgical removal leading to risks of injury to brain tissue. Biodegradable implantable sensors would eliminate the risks while providing sensing when needed. Here, we demonstrate a bioactive glass (BaG)-based hydration sensor. A fluorine (CaF_2_) containing BaG (BaG-F) was produced using a melting manufacturing technique. The structure and electrical properties of the resulting constructs were evaluated to understand the electrical behaviors of this BaG-based sensor. The synthetic process of producing the BaG-F-based sensor was validated by assessing the electrical properties. We demonstrated that this BaG-F chemical composition is highly sensitive to hydration, and that electrical activity (resistive-capacitive) is induced by hydration and reversed by dehydration. These properties make BaG-F suitable for use as a humidity sensor to monitor brain edema and consequently provide an alert for increasing intracranial pressure.

## 1. Introduction

Brain edema can develop following trauma and surgery, and can lead to increased intracranial pressure (ICP) and subsequent serious complications including death [1]. Therefore, a number of approaches for continuous intracranial monitoring have been investigated [2]. The use of implantable sensors can be useful; however, they need to be removed after recovery and this is associated with an increased risk of inflicting injury to the brain [3]. The development of biodegradable sensors for this purpose would offer a significant advantage obviating the need for a second procedure for device removal [4]. Biodegradable sensors are largely polymer based [5], however, the hydrophilicity of most polymers leads to swelling and water absorption, which can cause premature failure of sensors due to early degradation of internal components. In addition, polymer degradation is associated with inflammatory responses and chronic inflammation [6-8], driving a need for alternative materials for intracranial sensing.

Biodegradable metals have been explored as the basis for intracranial sensors, but the constraints on their use includes the accumulation of degradation by-products, which can have toxic effects [9]. Silicon (Si)-based materials can also be used in implantable sensors [10]; however, their degradation can produce reactive oxygen species that cause cell death [11]. Due to its slow degradation and dielectric properties, silicon oxide (SiO_2_) has been used mainly as a sensor encapsulating and interlayer dielectric, which has been used in active neural interfaces [12]. In this context, most studies on Si-based implantable sensors have demonstrated their effectiveness in detecting parameters such as pressure, temperature and pH [9]. In addition, Si-based biomaterials have the potential to contribute significantly to the development of new dielectric matrix concepts, mainly because the thermal, mechanical and optical properties of these amorphous structures are controllable [13]. However, bioceramics used in sensors generally have low reactivity, which is not recommended for use in monitoring the edema that develops after surgery or trauma because these sensors can also lead to an increase in ICP [14].

To address this need, we sought to develop a hydration monitoring sensor based on biodegradable Si-based BaG, which has favorable dissociation kinetics and reactivity for this clinical application, and when controlled association with chemical elements are used the physiological, mechanical and electrical effects can be modulated. In this context, electrical sensitivity of BaGs is; however, high, i.e. capacitive character and the speed electrical transfer mechanisms into material matrix can easily be modified even completely altered. This has been proven through impedance measurements [15], where magnesium-zinc-calcium (MgZnCa) alloy hybrid glass was used as a material for the control of hydrogen release from biodegradable implants [16]. It was demonstrated that the absence of dislocation-based plastic-deformation resulted in favorable strength and elasticity compared with those of a competing crystalline Mg derivative [17]. Recently, a humidity sensor with high sensitivity based on cellulose nanofiber was developed but this device is not biodegradable [18].

Electrochemical impedance measurements of these types of materials were seen to provide a quick, direct and sensitive approach for the detection of DNA sequences using a silicon (Si/SiO_2_) transducer with peptide nucleic acid (PNA) as the probe layer [19]. Other structural studies of the current flow mechanism within the vitreous borosilicate matrix prove that the electrical conductivity, measured by impedance spectroscopy, increases with the amount of B_2_O_3_ added [20]. The activation energy of fluoride, fluorophosphate and phosphate glasses are similar when derived by conductivity or electric relaxation, presenting some deviation at fluorophosphate-type glass. Also, the conduction phenomenon based on the ionic-polaron hopping with the hopping rate being thermally activated is shown for all of these glasses, whereas impedance plots and activation energy evolutions show that the electrical characteristics follow a non-Debye behavior [21].

To extend these observations and develop a biodegradable BaG sensor, we intercalated CaF_2_ into SiO_2_-Na_2_O-CaO-P_2_O_5_. CaF_2_ is an insulator that may be integrated into microelectronics, moreover, its presence in the glass matrix may improve the mechanical resistance of medical devices [22]. *In vitro* and *in vivo* biocompatibility testing of another type of BaG-F demonstrated an absence of inflammatory and toxic processes [23]. The chemical composition of our biomaterial may be controlled enabling customization for specific implantation sites, pathology type, and patient age and gender. CaF_2_ intercalation can, therefore, be useful for developing implants for use in elderly patients as it slows down the biodegradation process at the implant-body fluid interface [24,25]. In the current study, a novel BaG-based sensor was developed. To demonstrate how the chemical content of the glass materials affects density and electrical capacity, we evaluated several materials for biocompatibility properties. The electrical sensitivity of glass elements was found to be high and maliable, i.e. the capacitive character and the speed of the electrical transmission processes in the matrix of the material can easily be modified and even completely eliminated [26]. Although this amorphous material possesses a lower ductility, it shows an appropriate Young’s modulus with strengths nearly three times, and elastic limit nearly four times, higher than those of a crystalline Mg derivate. The novelty of the material used for developing these sensors is based on the controllability of its composition, fluorine content in particular. Furthermore, in the fabrication procedure, we did not dope our materials with fluorine, but we reached the stage of substitution at the network modifiers level while keeping the vitreous structure and bioresorbability limits. Regarding its electrical distinction, results obtained after the electrical impedance measurement in air ambient, BaG-F showed that the measured impedance can be modelled as a capacitor. It has a high sensitivity to changes in hydration (humidity) and; therefore, may be an exceptional candidate for monitoring brain edema, and help to intervene before a serious increase in ICP develops and thus prevent death.

## 2. Materials and methods

### 2.1. Electrical measurements

The dielectric measurement was realized through a laboratory measurement system based on an impedance analyzer and copper electrode. Fluorine bioactive glasses (BaG-F_x_) were measured although copper electrode that connected to the impedance analyzer (Agilent 4294A, Zeiss, Germany). Each side of the BaG-F specimen was attached to an electrode of the impedance analyzer (Figure 1). BaG-F impedance was measured in the frequency range 40 Hz to 30 MHz with amplitude of 500 Mv. The corresponding current was increasing with the frequency from some pA to 4 mA to better characterize the humidity impact on the electric behavior of BaG. To characterize the time dependence of impedance (Z), BaG-Fx were kept under an atmospheric pressure of 1 atm at an ambient temperature and humidity of 25°C and 30% RH respectively (Figure 1a). The RH environment was formed using saturated salt solution of potassium sulfate (K_2_SO_4_) 95 % RH (Figure 1b). The measurements were carried out under ambient conditions for three minutes.

**Figure 1.**
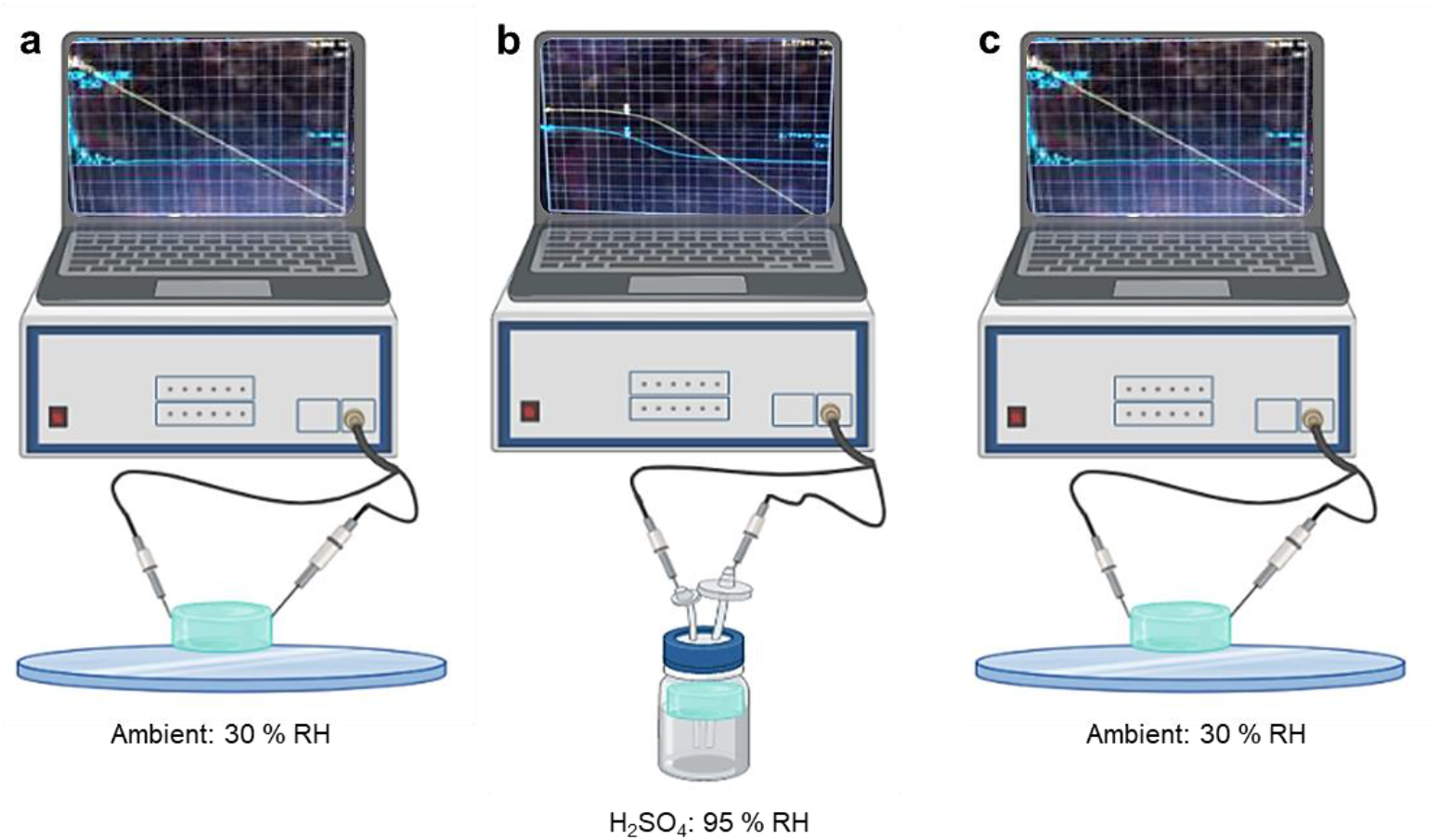
Measurement set-up for fluorine bioactive glass (BaG-F) based sensors showing the conductivity changes induced by humidity created by potassium sulfate solution (H_2_SO_4_). a) Capacitive character in ambient. b) Resistive-capacitive in high humidity. c) Reversal of capacitive character in ambient environment. The icons were taken from Biorender.com. Curves were derived from our study in the figure.

## 3. Results and discussion

Manufacturing of BaG-F-based sensors was performed by varying the weight content of CaF_2_ in BaG to maximize incorporation of fluoride (F^-^) in the glass network without destabilizing it and respecting the weight contents limits of each oxide to ensure bioactivity and biodegradability of all BaG-F_x_ [27].

### 3.1. Physical characterization of fluorine bioactive glass (BaG-F) based sensors

#### 3.1.1. Structural analysis

XRD demonstrated phase stabilization of the fabricated bioactive glasses (Figure 2).

**Figure 2.**
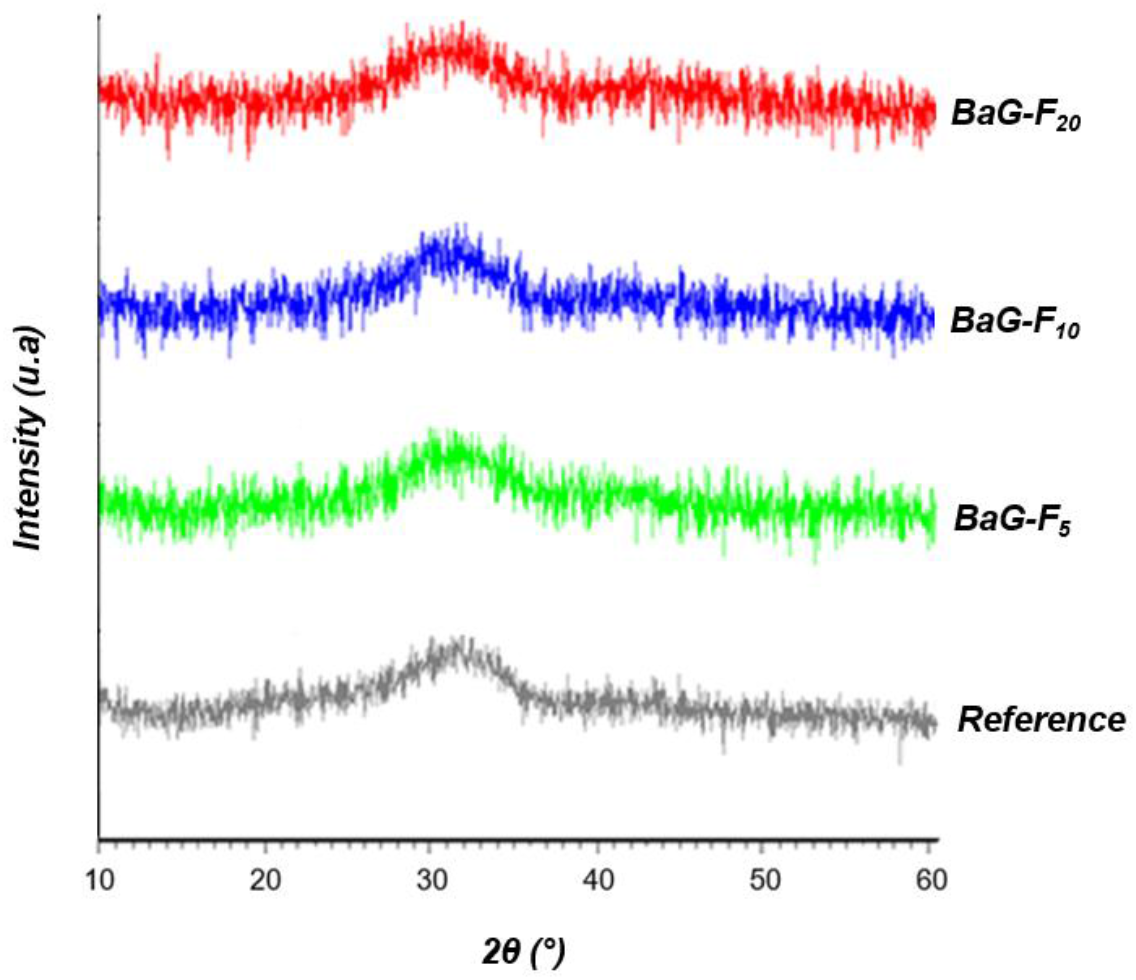
X-ray diffraction (XRD) patterns of fluorine bioactive glasses (BaG-F_x_).

This type of halo (24°< 2θ <38°) corresponds to the diffusion phenomenon in amorphous materials [20]. Therefore, vitreous and stable structure of all BaG-F_x_ was proven even after intercalation of CaF_2_ at a content of 20 wt.%. Identification of crystalline phases by X-ray diffraction was impossible for this type of material because of periodicity absence of atomic arrangement within vitreous network. No periodicity or stacking of planes at long distances, characteristic of pure glass. This promotes the phenomenon of electrical conductivity and fluid current flow through free electrons in purely disordered matrix which does not have boundaries of crystal reticular planes. Investigation of different molecular configurations of disordered glassy network of BaG-F_x_ was carried out (Figure 3).

**Figure 3.**
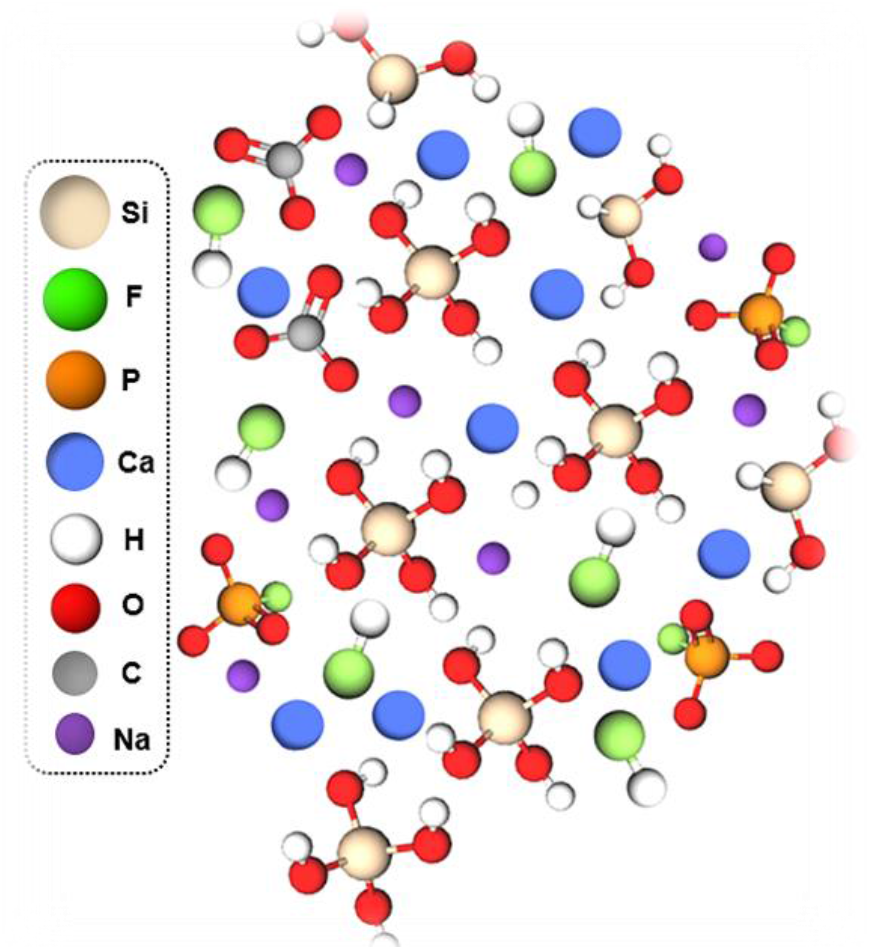
Investigated three-dimensionally (3D) structure of fluorine bioactive glasses (BaG-F_x_).

The electronegativity and electronic configuration of formers and modifiers elements (Table 1) may explain these molecular groups shape. The monatomic anion, fluoride (F^-^), with its important electronegativity, will establish changes in the electrical conductivity between the chemical bonds of BaG-F_x_.

**Table 1.** Physical and atomic properties of fluorine bioactive glasses (BaG-F_x_) components.

| *Chemical Element* | *Valence Layer* | *Electronegativity (Pauling scale)* | *Oxidation State* | *Electrical Conductivity (s.m<sup>-1</sup>)* |
| --- | --- | --- | --- | --- |
| <sup>14</sup>Si | 3s<sup>2</sup> 3p<sup>2</sup> | 1.9 | +1, +2, +3, +4 | 2.52 × 10<sup>-4</sup> |
| <sup>9</sup>F | 2s<sup>2</sup> 2p<sup>5</sup> | 3.98 | -1 | - |
| <sup>8</sup>O | 2s<sup>2</sup> 2p<sup>4</sup> | 3.44 | -2, -1 | - |
| <sup>20</sup>Ca | 4s<sup>2</sup> | 1 | +2 | 29.8 × 10<sup>6</sup> |
| <sup>11</sup>Na | 3s<sup>1</sup> | 0.93 | +1 | 21 × 10<sup>6</sup> |
| <sup>15</sup>P | 3s<sup>2</sup> 3p<sup>3</sup> | 2.19 | ± 3, 5, 4 | 1.0 × 10<sup>-9</sup> |

Understanding the atomic properties of each constituent element of this matrix type is important [28] to better explain the chemical bonding alterations that can affect the glasses morphology and electrical behavior. Table 1 illustrates the atomic properties of each chemical element constituting BaG-F network.

### 3.2. Electrical behavior of fluorine bioactive glass (BaG-F) based sensors

In this investigation, we assume that the measurement area is the same for all BaG-F_x_. The measuring device usually consists of connecting the material (BaGs) to be analyzed to a resistance, inductance and capacitor (RLC) meter or impedance analyzer. This technique is a simple, fast and non-destructive analysis that provides data on dielectric properties of new materials subjected to an electric flow. All BaG-F_x_ were analyzed over a wide frequency range from 50 Hz to 30 MHz, generated at a constant applied voltage of 500 mV (peak-to-peak). The frequency range included lower frequencies in order to properly select sensitivity of this new type of material. For all BaG-F_x_ samples, capacitive character in the ambient air is detected. The measured impedance can then be modeled as a capacitor (C) (Figure 4a). The impedance (Z) can be calculated from the expression for the modulus in equation 1:

**Figure 4.**
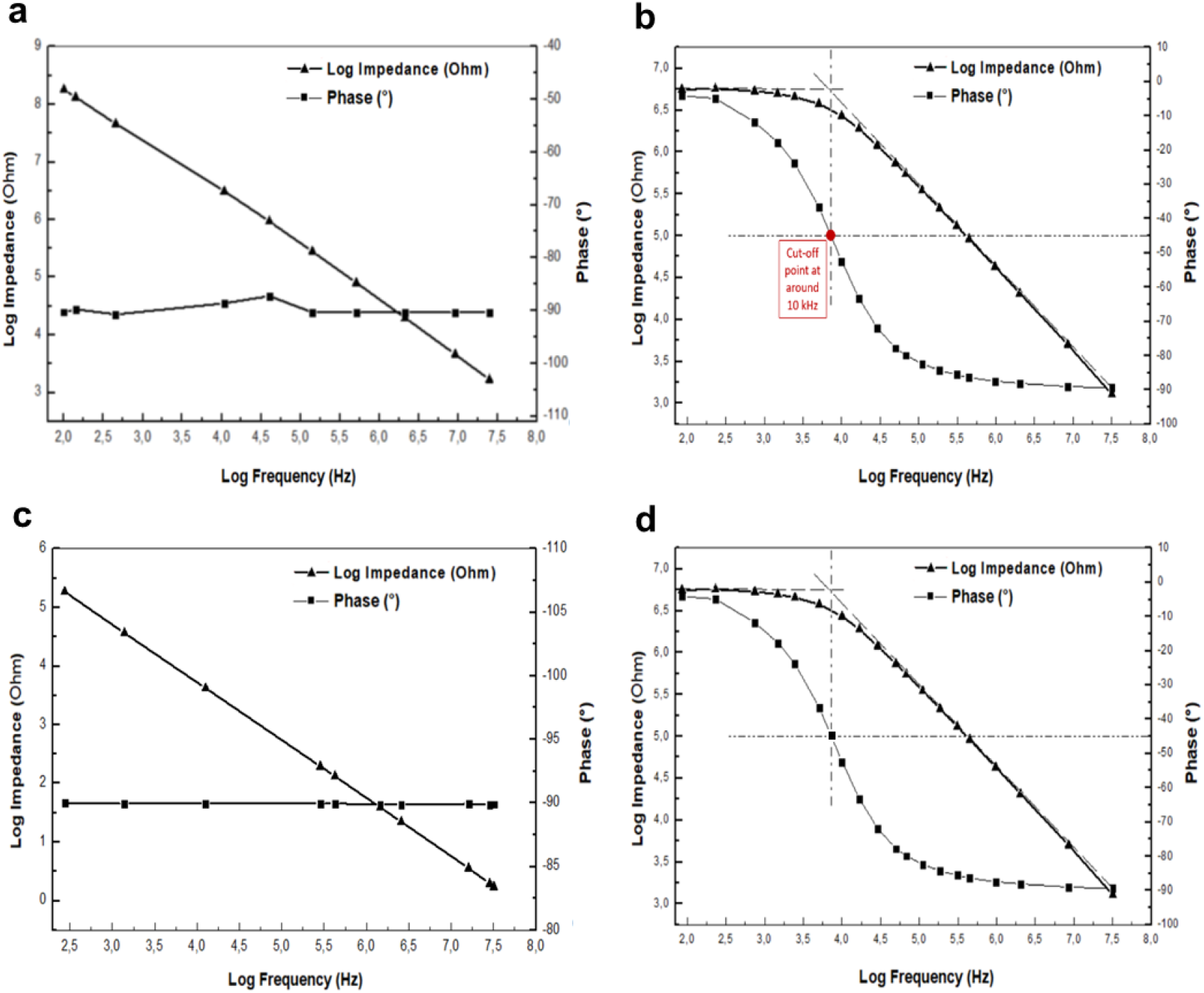
Reversible resistive-capacitive behavior for bioactive glass-based sensor (BaG-F_10_). a) Capacitive character in ambient. b) Resistive-capacitive character in humidity (95 %RH). c) capacitive character returns within 5 seconds after removal of BaG-F_10_ from the humid medium d) Reversible resistive-capacitive character after BaG-F_10_ further contact with humidity.

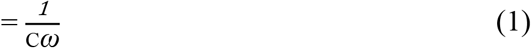

In this process, free movement of atomic electrons on the BaG-F_x_ surface ensures the flowing ionic charge transfer, especially since our BaG network was based on a semi-conducting chemical element which is silicon (Si). Moreover, fluoride (F^-^) is at the top right of chemical periodic table being the most electronegative atom (Table 1). It has a great tendency to extract itself from valence layer by bonding with another free electron reaching the surface. Consequently, presence of a reactive element (F^-^) insures a perfect electrical charge succession between network fragments of BaG-F_x_. In addition, energy provided by electric flow is sufficient for the decomposition of CaF_2_ by losing one fluoride atom per molecule, this desorption at the Si/CaF_2_ interface is manifested by breaking of Si-F bonds creating Si-Ca which promotes electroconductivity by calcium cation (Ca^2+^).

On the other side, measurements carried out in the ambient air for BaG-F_x_ show that in capacitive developments the phase is constant and relatively fixed at -90°. This outcome is a logical result, as such negative phases are usually associated with capacitive performance. Besides, it can be stated that positive phase shifts can be obtained by inductive components [29]. Generally, bioactive electronic systems have to meet high sensitivity to physiological changes (pH, water presence, temperature). In addition, special methods are needed for the manufacture of medical devices with delicate microstructure, while avoiding hydrolytic disintegration of the materials during fabrication process [30]. In this context, the focus was on studying the effect of humidity on the electrical characteristics of BaG-F_x_.

With the same electrical parameters as measurements made in ambient air, rapid change in electrical behavior of BaG-F_x_ at 95% RH is interesting and encouraging to further explore their response to water content. Precious electrical behavior has been observed for this type of biomaterial, for example BaG-F_10_ (with average medium CaF_2_ added amount, Figure 4b).

In fact, BaG-F_10_ showed two behaviors: before the cut-off point, the resistive character predominates. Whereas for frequencies above this critical frequency point, which corresponds to approximately the -45° phase value, material capacitance power was high, and the capacitive aspect persists. It has been proven that apparent resistance and capacitance of materials strongly influence magnitude and phase of low frequencies [31]. In the frequency range before cut-off point, impedance (Z) of BaG-F_x_ can be equated to a resistance (R), while capacitive effects can be neglected before the cut-off point. The biomaterials have resistive character, and the phase is zero. Therefore, Z is equal to R. For the frequency range above 10KHz, Z can be represented as an equivalent circuit representing the capacitance (C) in parallel with the resistance (R). R and C are generated by humidity measuring device. In this range, the impedance (Z) can be determined using equation 2:

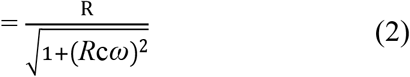

This experiment proves that whatever the measurement duration of 3 min or 30 min, BaG-F_10_ has a resistive and then a capacitive character (depending on the frequency range) and if it is removed from the humidity, i.e. water molecules (H_2_O) presence, it will quickly get free of adsorbed water and it will return to its initial capacitive character within 5 seconds (Figure 4c). This reversible behavior (Figure 4a-d) is due to high reactivity of BaG-F_x_ versus humidity.

The same behavior was seen with other BaG-F_x_ glasses. This study adapts the hypothesis that surface contact of BaG-F_x_ has been affected by chemical bond type changes and molecular reorganizations. Through physical adsorption mechanism, the charge exchange between H_2_O and BaG-F_x_ is well performed. From the first point of view, chemical contact in H_2_O/BaG-F_x_ surface is known as a hydrolysis reaction [32]. SiO_2_ is the main BaG forming compound which is configured as a vitreous network with tetrahedral units and the four oxygen atoms in silicate tetrahedron (SiO^-^_4_) are shared [20]. By hydrolysis, Si-O bonds are typically broken (step 1 in Figure 8). Depolymerization results in the formation of orthosilicates or the silanol group Si(OH)_4_ [33]. Aggregation of these groups forms a three-dimensional (3D) layer that acts as an insulating screen for the current (step 2 in Figure 5), giving the biomaterial a resistive character in the frequency range below the 10 kHz cut-off point (Figure 4b). Moreover, Si(OH)_4_ hydroxide network acts as an insulating block due to its steric hindrance which limits the electrical conductivity in the low frequency range. However, these silicon hydroxides are not stable until polymerization by liberation of H_2_O ions and then dehydration to return to the mis-ionic Si-O-Si bond (step 3 in Figure 5) which is the active monomer that assures a smooth electrical transfer in the high frequency range. In addition, the high electronegativity of oxygen (O) and silicon (Si) (Table 1) favors current flow through this Si-O-Si siloxane bridge (Figure 5).

**Figure 5.**
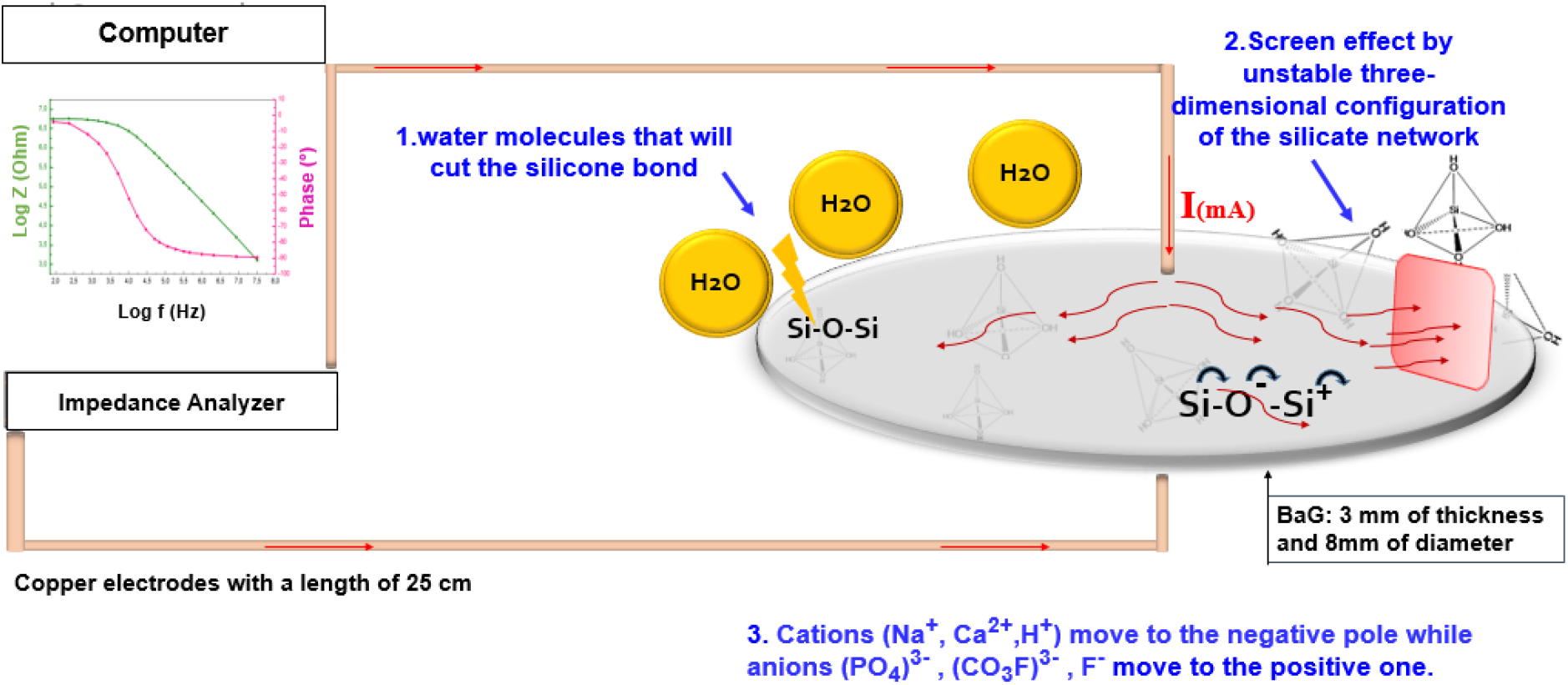
Illustration of the physicochemical mechanism behind the resistive-capacitive behavior of biomaterials.

From the second point of view, F^-^ plays an important role in these structural changes upon BaG contact with water. Recent research proves a reversible decrease in voltage as a function of the content of CaF_2_ added to the silicate network (SiO_2_) through electrical measurements on the CaF_2_/SiO_2_ interface of ionic/covalent bond respectively [34]. This process results from substitution of CaF_2_ by CaO at the interface. CaF_2,_ being very reactive during electrical bombardment, provides the dioxygen (O_2_) for the exchange to take place. This allows to highlight hysteresis phenomena and negative differential resistance behavior for fluorine-silicate material. These dynamic effects could be modelled using an equivalent electrical circuit made of discrete elements (resistors-capacitors as in accord with our results).

## 4. Conclusion

It is possible to fabricate fluorine containing bioactive glasses that are characterized by stability, high density and electrical conductivity. It is possible to have hydration-induced electrical activity behavior of the BaG (becoming resistive and then conductive), which can be reversed upon dehydration. Because of these structural, chemical and electrical properties of fluorine-implantable devices, without losing their mechanical and biological assets, these chemically reactive BaGs can serve as biosensors. This is the first study to demonstrate the potential of BaG-F to serve as a sensor for the monitoring of hydration and humidity. Our results, therefore may provide an exceptional candidate for monitoring of intracranial edema and address an important clinical problem. Further simulation studies are in progress to optimize this sensor further (matching conductivity and humidity). In future, they will have potential applications in monitoring of clinically relevant edema such as that of the brain.

